# Passive Acoustic Dynamic Differentiation and Mapping: A Time-Domain Passive Cavitation Localization and Classification Approach

**DOI:** 10.1101/2025.04.08.647829

**Authors:** Nathan Caso, Krunal Patel, Tao Sun

## Abstract

1

**Objective:** Passive cavitation imaging and control has explored various beamforming algorithms to balance resolution, imaging artifacts, and computational speed. Optimizing these parameters is essential for clinical translation, as precise cavitation localization and dosage control are critical for Focused Ultrasound (FUS)-based targeted therapies, minimizing unintended tissue damage. Among commonly used methods, Delay-Sum-Integrate (DSI) and Robust Capon Beamforming (RCB) have demonstrated effectiveness but are limited by either significant artifacts or the need for extensive parameter tuning.

**Methods:** This work introduces Passive Acoustic Dynamic Differentiation and Mapping (PADAM), which adapts the Multiple Signal Classification algorithm to the time domain to improve cavitation localization.

**Results:** PADAM achieves up to a 6-fold improvement in lateral beam-width compared to RCB, and a 4-fold reduction in mean-square intensity of artifacts. It further unveils a novel physical insight: its input parameter dynamically gauges the richness of an incoming signal’s frequency content. This feature enables a more physically defined and intuitive parameter for distinguishing between stable and inertial cavitation based on spectral characteristics, simplifying parameter selection and enhancing the framework for cavitation monitoring and control.

**Conclusion:** With its ability to improve resolution, reduce artifacts, and provide computational efficiency, PADAM represents a promising advancement for precise cavitation localization and therapy monitoring.

**Significance:** This work introduces PADAM, a novel time-domain passive cavitation imaging method that offers superior resolution and artifact reduction compared to DSI and RCB. Its physically intuitive input parameter enables dynamic differentiation between stable and inertial cavitation, enhancing precision in the monitoring and control of FUS therapy.

## 2 Introduction

Passive Cavitation Imaging is an emerging field within ultrasound imaging and signal processing, gaining significant attention for its role in monitoring cavitation facilitated therapeutic ultrasound applications including targeted drug delivery, focal ablation, and others. These therapeutic applications often utilize a focused ultrasound (FUS) transducer to induce acoustic cavitation with or without seeded bubbles, enabling spatially targeted therapeutic effects. For instance, cavitation can facilitate the release of microbubble-encapsulated drugs, allowing treatments to reach regions that are typically inaccessible to conventional therapies and the immune response. This is seen in hypoxic ischemic areas in tumors and across the blood-brain barrier [1] [2]. Both preclinical and clinical studies have demonstrated enhanced therapeutic efficacy when cavitation-based approaches are applied to conditions such as Alzheimer’s disease, glioblastoma, and other pathologies [3] [4] [5] [6] [7] [8] [9].

However, the stronger mode of cavitation, commonly referred to as inertial cavitation, can be highly destructive, leading to undesired cell damage and death due to mechanical stress and hyperthermia [5] [10]. Non-invasive control and monitoring of cavitation-facilitated FUS therapy remain as a key challenge, traditionally addressed using MRI-based techniques or passive acoustic imaging, the latter being more cost-effective and time efficient [11] [12] [13]. Passive acoustic mapping (PAM) utilizes a passive listening device to estimate cavitation power and localize the treatment, making it a topic of significant interest in array signal processing [14]. High-resolution beamforming algorithms are crucial for improving the clinical viability of this passive imaging. Conventional approaches, such as Delay-Sum Integrate (DSI) and the Robust Capon Beamformer (RCB), have been used for cavitation power estimation and localization, and each display inherent trade-offs in resolution, speed, and artifact suppression [15] [16] [17] [18] [14] [19].

### Classical Passive Beamforming Techniques

The Delay-And-Sum (DAS) beamformer remains a commonly used method due to its computational efficiency [15] [16] [17] [18]. DAS estimates pixel intensity by summing delayed signals from multiple transducer channels based on their respective distances to the pixel location [20]. A more advanced variant, Delay-Sum Integrate (DSI), extends DAS to cases where the incoming wavefront’s time is not known. By integrating across all available time indices, DSI provides a long-exposure-like reconstruction of cavitation events [21]. DSI may be implemented in either the time or frequency domain, with accuracy and speed improvements achieved by selecting frequencies associated with stable or inertial cavitation [22]. Although DSI is well-suited for real-time applications due to its computational efficiency [23] [24] [25] [26], it suffers from comparatively poor resolution and a characteristic tail artifact, a nonphysical effect resulting from the overlapping wavefront delays across channels.

To mitigate these limitations, adaptive beamformers have been explored, notably the Capon Beamformers (CBs) and their enhanced version that prevent self-nulling, the Robust Capon Beamformers (RCB) [14] [19] [23]. Several studies have shown that RCB improves localization accuracy compared to DSI, aligning well with measured bio-effects in vitro [22]. However, despite its advantages in resolution, RCB is computationally intensive and frustratingly requires fine-tuning of a steering vector uncertainty parameter, *ε* [14]. This parameter, defined as:

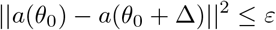

quantifies the allowable uncertainty in the steering vector [27]. Selecting an optimal *ε* remains a challenge, as useful values can vary by orders of magnitude. Some works suggest tuning it to around 8.45 [19]. We noted values between 1 × 10^*−*9^ and 0.3 generated images the with greatest tail-artifact suppression for the experimental setup. Eigenspace-based RCB has been proposed to reduce the variance of the input parameter [28], but *ε* remains an empirically determined factor rather than a physically measurable parameter, limiting its practical implementation.

### Proposed PADAM Approach

To address the limitations of existing beamformers, this work introduces the Multiple Signal Classification (MUSIC) direction-finding beamformer to the field of passive cavitation imaging. MUSIC is established in far-field array sensing (such as radar and sonar applications) for high-resolution and high signal to noise ratio (SNR) direction-finding, yet an adaptation to passive cavitation imaging remains largely unexplored. This study evaluates the feasibility of time-domain MUSIC as a solution to the trade-offs between resolution, speed, and artifact suppression observed in existing beamformers. The remainder of this work is organized as follows: Section 3 is a Methodology that includes 1) a derivation of the MUSIC signal model, 2) an explanation of MUSIC in the frequency domain, 3) a derivation of MUSIC in the time-domain, 4) an explanation of the computational models used, 5) an explanation of the experimental methods used, and 6) an explanation of the metrics used to compare the beamformers. Section 4 (Results) presents findings from both in-silico and in-vitro experiments, analyzed using the aforementioned metrics. Section 5 (Discussion) compares PADAM’s capabilities and limitations against DSI and RCB. Section 6 (Conclusion) summarizes key findings and suggests directions for future research.

## 3 Methods

### MUSIC Signal Model

The original MUSIC is a frequency domain direction-of-arrival beamformer that uses Eigenvalue analysis and assumes the presence of *m* coherent scatterers [29]. It divides the received radio frequency (RF) data into a noise and signal subspace, and assumes the incoming signal can be modeled as the summation of discrete sinusoidal sources and noise.

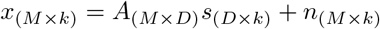

Here, *A* as the matrix of steering vectors corresponding to *M* channels with *D* actual sources, *s* is the signal vector for a snapshot with *k* samples, and *n* is the noise vector. For convenience, we will use *k* = 1 for the remainder of the signal model. Next, we define the spatial covariance matrix *R*_*xx*_ as the expected value of the sample covariance matrix:

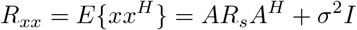

Where the signal covariance matrix *R*_*s*_ = *E* {*ss*^*H*^} is of size *D* × *D*, and the spatial covariance matrix is of size *M*× *M*. To satisfy the assumption that *R*_*xx*_ = *E* {*xx*^*H*^}, multiple snapshots are often used:

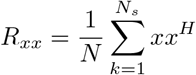

Because *R*_*xx*_ = *AR*_*s*_*A*^*H*^ + *σ*^2^*I*, the rank of *R*_*s*_ is equal to the number of sources *D*, and the rank of the noise matrix *σ*^2^*I* is zero, the rank of *R*_*x*_*x* is also *D*. This means that there are *D* nonzero Eigenvalues of *R*_*xx*_ corresponding to the signal, and *M* − *D* near-zero Eigenvalues corresponding to noise. The Eigenvalue decomposition of *R*_*xx*_ is:

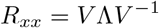

Where 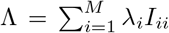 and *λ*_*i*_ is the i’th Eigenvalue, and *V* is a matrix of the Eigenvectors, sorted in the order of the largest to smallest Eigenvalues.

### Classical MUSIC Reconstruction

With the signal model established, reconstruction using MUSIC proceeds as follows: The incoming signal is either accumulated or divided into “snapshots”, *x*_(*n*_*t×nc*)*i*, which are averaged in the frequency-domain. *n*_*t*_ is the number of time samples, and *n*_*c*_ is the number of transducer channels, so *x* is a matrix with *n*_*t*_ rows and *n*_*c*_ columns. The spatial covariance matrix estimate is also generated during this step by taking the inner product:

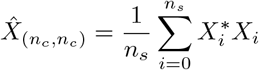

Where *n*_*s*_ is the number of snapshots used. Next, the Eigenvalue decomposition of the averaged spatial covariance matrix estimate is computed:

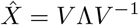

From this decomposition, the number *m* of scatterers is selected by the user, assuming the signal rank *D* of 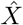 The *m* largest Eigenvalues are selected from the decomposition, where *m < M*.

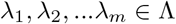

The associated Eigenvectors from each of these *m* Eigenvalues form the signal subspace, *V*_*s*_ ∈ (*M* × *m*), and the remaining *n*_*c*_ − *m* + 1 Eigenvectors form the noise subspace, *V*_*n*_ ∈ (*M* × *M* − *m*). For each direction (or pixel location in the near-field), the pseudo-spectral intensity is calculated as the inverted inner product of the steered noise subspace:

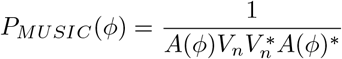

Notably, from this derivation, MUSIC does not yield a power estimate. It is instead inversely proportional to the noise power; when the noise subspace is orthogonal and uncorrelated (i.e., the signal subspace is highly correlated), then the MUSIC pseudo-spectrum is large.

### PADAM - MUSIC Time-Domain

In this work, we adapt the MUSIC algorithm for the time-domain to form the PADAM algorithm. The input signal *x*_(*n*_*t,nc*) is delayed in time according to the geometric distance between each transducer’s position *p*(*x, y*) and a pixel location *a*(*x, y*), using the one-way time of flight at the speed of sound, following the DSI approach:

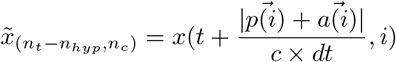

Where *i* is the index of a channel or transducer. The delays may be computed using a circular shift including all samples, or *n*_*hyp*_ samples corresponding to the hyperbola shape of the delay across channels may be omitted, as done in this work.

Next, the delayed signal is treated as *n*_*t*_ − *n*_*hyp*_ snapshots in time, and averaged in the timedomain:

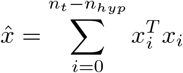

As in the frequency-domain approach, the Eigen analysis is performed, but in this case, the entire decomposition is real-valued:

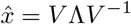

*x*The pixel value is calculated from the noise subspace as before, but because a delay has already been applied, the subspace inner product is no longer steered during this step. Specifically, the steering vector *A*(*ϕ*) is replaced with a vector of ones, which is equivalent to the two-dimensional summation of the matrix elements:

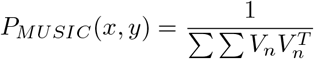

The algorithm is laid out step-by-step in Algorithm 1

#### Algorithm 1

PADAM Time-Domain

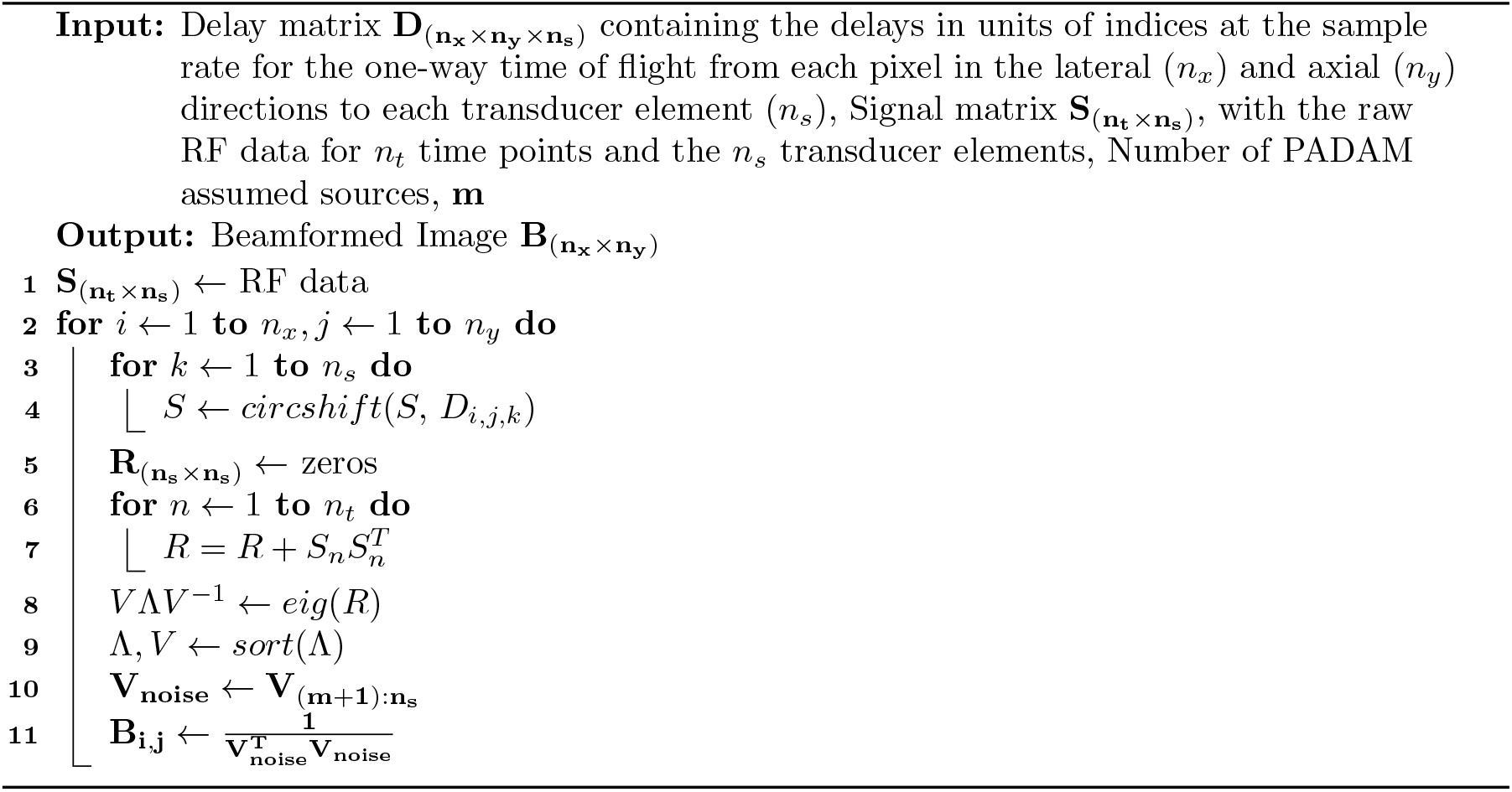

In comparison to classical MUSIC, our PADAM time-domain approach offers the advantage of more extensive snapshot averaging, at the sacrifice of a potentially expensive average calculation and the need to recalculate the Eigenvalue decomposition for each pixel. One significant advantage of this time-domain method is its simplicity. The frequency domain method requires careful management of snapshot lengths to retain relevant frequency information when splitting a frame of RF data into snapshots. Alternatively, averaging multiple frames of RF data to improve signal-to-noise ratio can come at the expense of temporal resolution. Moreover, the time-domain method eliminates the need for common signal processing steps, such as windowing and overlapping, needed for frequency domain averaging, offering a more straightforward approach without sacrificing essential data integrity

### Computational Model

Vokurka’s bubble signal model provides a computational framework for simulating the time-series behavior of cavitating microbubbles using stochastic distributions in several aspects. This model offers a unique platform for testing beamformer performance, as the location of a bubble source can be arbitrarily controlled in both space and time. The model is based on Vokurka’s observations on cavitation behavior, where a cavitation event can be modeled over time by the equation [30] [31] [32]:

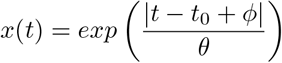

Where *t*_0_ represents the time of the cavitation event, *ϕ* is a stochastic variable representing a small random time shift, and *θ*, referred to by Vokurka as the “time-condition”, controls the width of the cavitation event’s build-up and ramp-down. *θ* can also be manipulated stochastically. The process may be repeated at regular time periods *T*, corresponding to a FUS pressure cycle’s period, and for many point-scatterers:

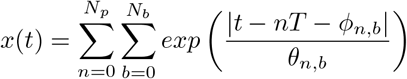

Where *N*_*p*_ is the number of periods, and *N*_*b*_ is the number of scatterers. To model the RF data received at a transducer’s location, *x*(*t*) is delayed by the propagation time between the scatterer’s location and the transducer, and scaled by 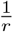 to account for proportional spherical spreading. Additionally, a bandpass filter is applied to mimic the frequency response of an imaging probe. The virtual probe used in this model consists of a 64-element linear array designed to simulate a Verasonics linear probe, with a center frequency of 3.21 MHz and a pass-band ranging from 1.2 to 5.2 MHz.

### Experimental Methods and Materials

To validate the computational results and compare beamformer performance for cavitation, an in-vitro model was used. In-house prepared lipid microbubbles were diluted and placed into a 2-mL Eppendorf tube. A 500-kHz FUS transducer (FUS Instruments, Toronto, ON, Canada) was positioned at its 2.5 mm focal length at a fr,equency of 0.5 MHz below the upright tube. A P4-1 imaging probe (Philips/ATL, Cambridge, USA, MA) was positioned flat on the radial axis of the tube at distances between 25 and 55 mm.

Cavitation was induced by the FUS transducer at an amplitude of 0.5 MPa for 10 seconds, using a 10 ms pulse length and a repetition frequency of 0.25 Hz. The P4-1 probe was also used to transmit a simple B-mode flash sequence for localization, followed by passive RF data acquisition for cavitation imaging. The B-mode and cavitation images were captured with a Verasonics Vantage 256 (Verasonics, Kirkland, WA, USA) system and beamformed on a Dell Precision workstation (Dell Inc, Round Rock, TX, USA) with an Intel Xeon W-2255 processor (Intel, Santa Clara, CA, USA).

To compare the relative performance of each beamformer with a skull phantom, a rat skull was positioned in a water tank 25mm below the FUS transducer. An L12-5 (38mm) probe (Philips/ATL, Cambridge, USA, MA) was positioned at a 90-degree orientation to the FUS focal axis, parallel to the anterior-posterior axis of the skull. The skull was rotated 45 degrees towards the L12-5 probe, so that both the FUS and any cavitation emissions would pass through parietal bone. A 1.2 mm hole was drilled in the skull in the eye cavity of the frontal skull bone, and a 1 mm inner-diameter PTFE tube (McMaster-Carr, Princeton, USA, NJ) was passed through the cranial cavity. In-house prepared lipid microbubbles were diluted and pumped through the tubing using a 5-mL syringe with an 18-guage tip. Cavitation was induced using a pressure of 1.5 MPa at 500-kHz for 10 seconds with 10 ms pulse lengths, and a pulse repetition frequency of 10 Hz.

### Comparison Metrics

To effectively compare the performance of the beamformers, several metrics were used. The point spread function (PSF) of a beamformer is the ideal metric, although it is spatially dependent: every pixel exhibits a slightly different point-spread. For this work, the full-width at half-maximum (FWHM) was used as the measure of beam profile width in both the axial and lateral directions.

The mean-square intensity (MSI) of the pixels serves as a measure of the amount of relative noise or cumulative artifact presence in an image. This metric encompasses both signal and artifacts; however, since the size of cavitating microbubbles is on the micrometer scale, an order of magnitude smaller than the millimeter scale used in ultrasound imaging applications, the cavitation source is point-like and much smaller than a single pixel. For the images generated in this work, a single, stationary cavitation source (such as those simulated by Vokurka’s model) would produce minimal signal intensity, while the overwhelming majority of observed pixel intensity comes from artifacts and the point-spread function. Therefore, MSI can be used to quantify artifacts and the effects of point-spread. MSI is calculated by:

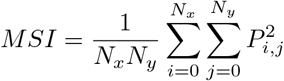

## 4 Results

### Localization and Artifact Reduction

To compare PADAM to RCB and DSI, the in-silico Vokurka model was used with a single cavitation model source positioned axially at 130 mm and centered laterally in a virtual probe’s field of view. The images were beamformed, and the output images were normalized on a linear scale for comparison, as PADAM is not a power-based beamformer and the intensity of the images cannot be compared directly. Figure 2 presents a comparison between DSI, RCB, and PADAM for this single source, where the number of assumed sources for PADAM is *m* = 1. PADAM exhibits a point-like source stretched by the axial resolution of the system, without a tail artifact, whereas RCB displays a much larger area without a tail, and DSI shows the classical X-shaped tail artifact.

**Figure 1:**
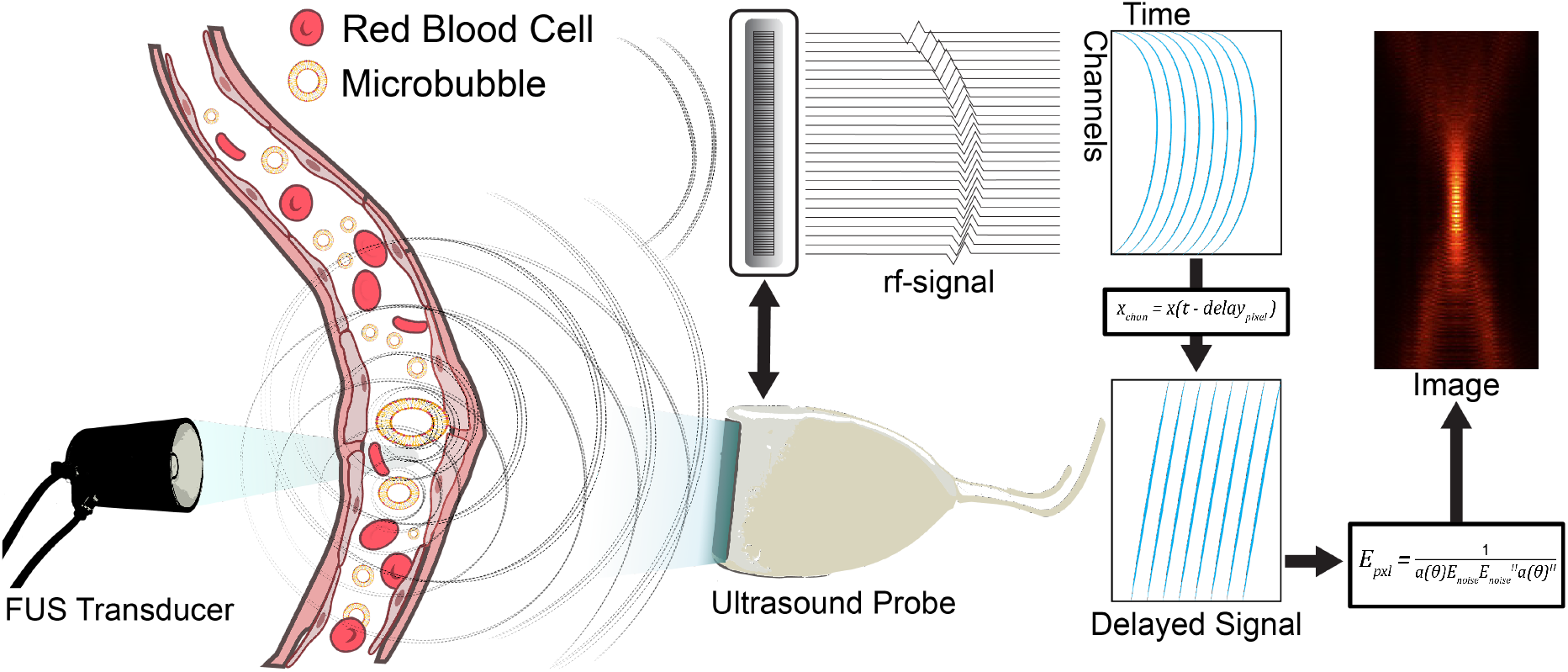
An illustration of the cavitation imaging setup, with a focused ultrasound generating a signal that cavitates lipid nanoparticles in the targeted vasculature. This signal is picked up by a traditional ultrasound probe in passive listening mode, and beamformed to generate cavitation images.

**Figure 2:**
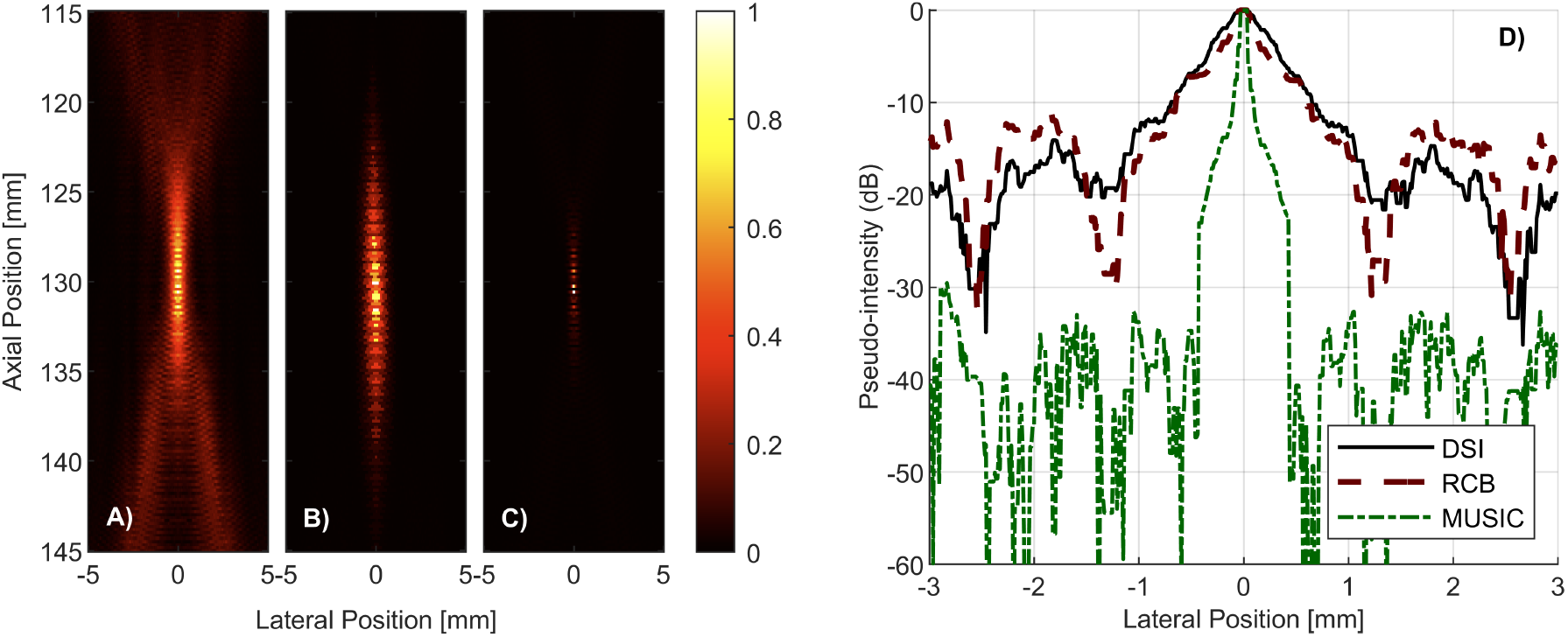
In-silico passive cavitation image using the Vokurka Model. A) depicts the passive cavitation image beamformed using Delay-Sum-Integrate. B) shows the same RF data beamformed using the Robust Capon beamformer, and C) shows the image beamformed using MUSIC with *m* = 1. Each image is normalized between 0 and 1, where the pixel values represent pseudo-power. D) Shows the axial beamwidth profiles for the three beamformers

The axial beam-widths for each beamformer are also shown in Figure 2, with PADAM demonstrating a significantly narrower width. These beam-widths are also summarized in Table 1.

**Table 1:** Axial Beam Widths (mm)

|  | DSI | RCB | MUSIC |
| --- | --- | --- | --- |
| -3 dB | 0.493 | 0.253 | 0.072 |
| -6 dB | 0.794 | 0.625 | 0.096 |
| -10 dB | 1.347 | 1.251 | 0.192 |

The Vokurka model was further used to quantify resolvability, as done in Figure 3, where two bubble sources initially superimposed at the same position are incrementally separated in the lateral direction. A 1-Dimensional axial image (or A-line) was constructed at the depth of the bubble for each bubble spacing, and these images are concatenated. To mitigate randomness in the stochastic measurements of the Vokurka model, 30 line-images were averaged at each source spacing. As seen in Figure 3, the RCB beam width appears sharper when the sources are spaced farther apart, but a visible artifact remains in the center. This artifact is also observed with DSI. Using the definition of resolution as the minimum distance at which two point sources remain distinguishable, the observed resolutions observed are approximately 1 mm for DSI, 0.65 mm for RCB, and 0.5 mm for PADAM.

**Figure 3:**
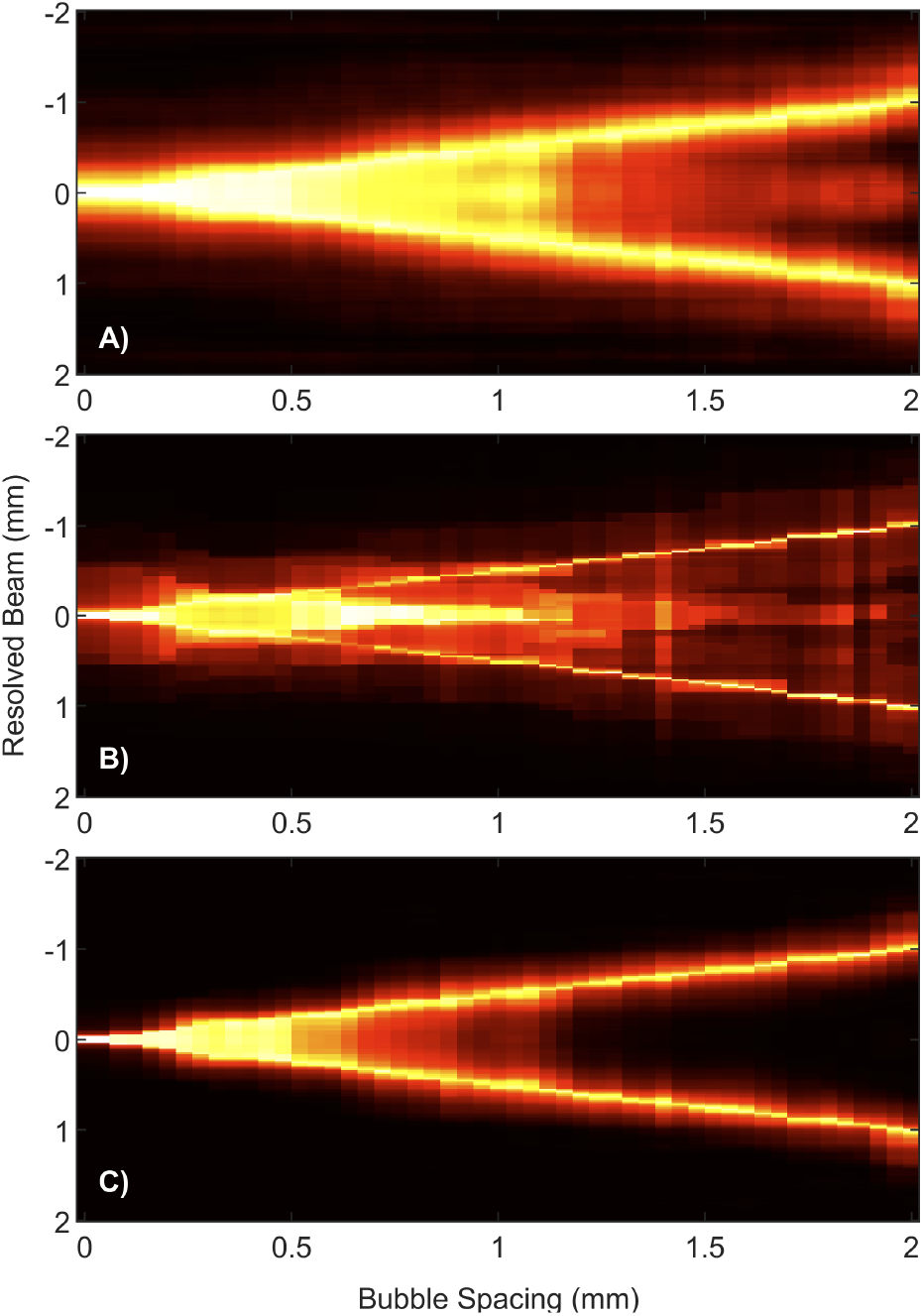
Point-source resolvability, where A) is DSI, B) shows RCB, and C) depicts PADAM with *m* = 1

To simulate the bubble cloud in the therapeutic focal target, we further placed the Vokurka modeled bubbles at random locations within an ellipsoid shape mimicking a FUS focal region. Clusters of bubbles serve as ideal test cases for artifact removal, as the tail artifacts between bubbles can merge and produce compounding effects in regions devoid of bubbles.

As seen in Figure 4 PADAM, with *m* = 1, exhibits significantly lower MSIs in comparison to DSI and RCB, largely due to its ability to suppress the tail artifact. RCB demonstrates a limitation wherein it exhibits a variable noise floor, contributing to its high MSI, despite superior suppression of the tail artifact relative to DSI. This issue appears to stem from the static *ε* coefficient. These findings indicate that PADAM is highly effective in noise reduction. However, as seen in Figure 4, PADAM tends to over-suppress the weaker point sources, an effect discussed in the following section.

**Figure 4:**
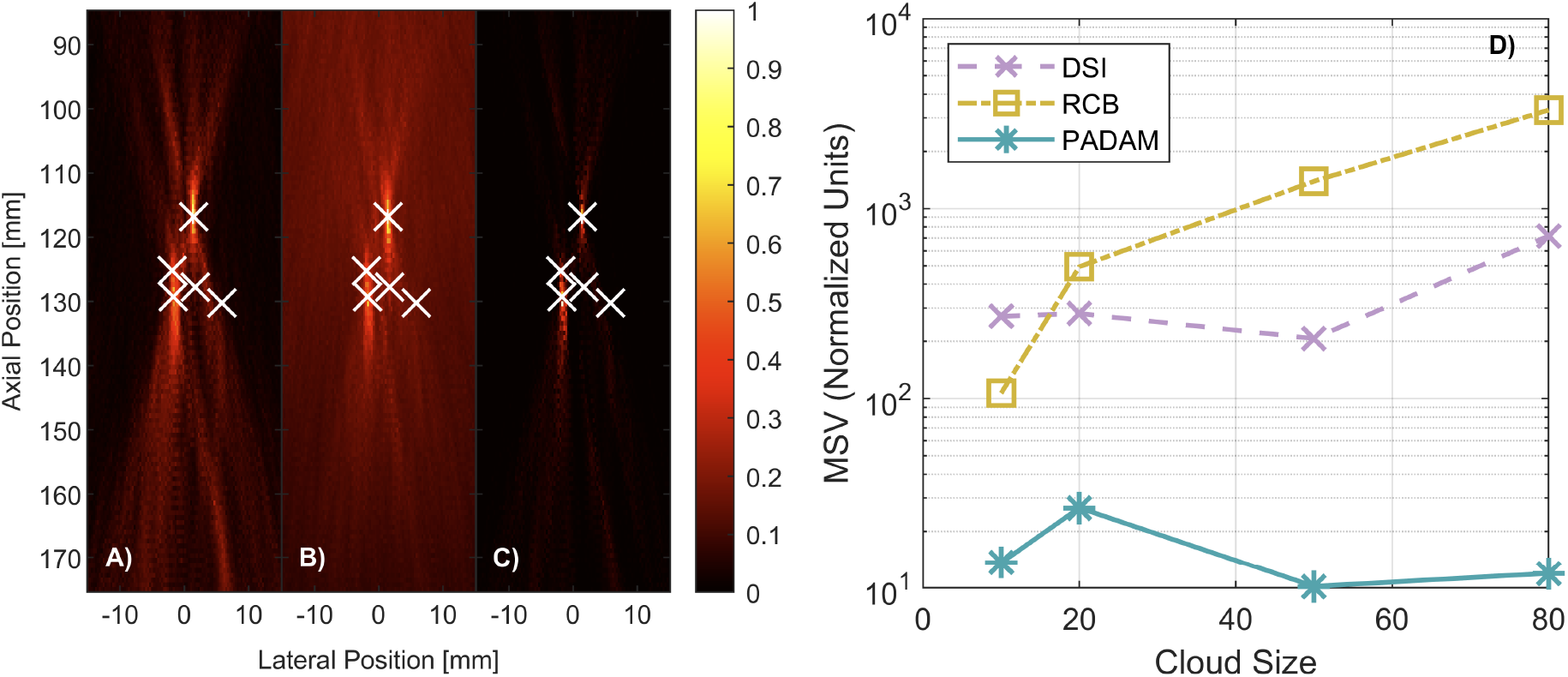
A representative image of a 5-source cavitation image, with each beamformer, and the exact source locations overlaid in white x’s. A) depicts DSI, B) depicts the same RF data beamformed with RCB, and C) beamformed with MUSIC. As in other figures, the images are normalized. D) depicts the MSI for each beamformer for four bubble cluster sizes: 10, 20, 50, and 80

### The PADAM Parameter *m*

As outlined in Section 3, PADAM operates by assuming a number of sources *m*, from which the signal and noise subspaces are determined using accepted and rejected Eigenvectors. This means that at each pixel, only the Eigenvectors corresponding to the *m* largest Eigenvalues are included. The largest *m* Eigenvectors correspond to the most correlated sources in the spatial covariance matrix at that pixel, effectively filtering out uncorrelated signals. Additional Eigenvalues in the spatial covariance matrix may represent “weaker” sources at other locations with lower correlation, while near-zero Eigenvalues primarily correspond to noise. Selecting the largest Eigenvalues ensures that the most correlated sources at a given pixel are passed through the beamformer.

In this line of thoughts, increasing *m* to larger integers may introduce leakage signal from other sources, as well as potential noise. A higher *m* allows weaker or less-correlated source to become visible in the image. Figure 5 displays a further confirmation of this concept, where a PADAM image is beamformed using a sine-wave with frequency *f* = 3.21*MHz* and amplitude 1 MPa and compared to a Vokurka source with a mean peak amplitude of 1 MPa.

**Figure 5:**
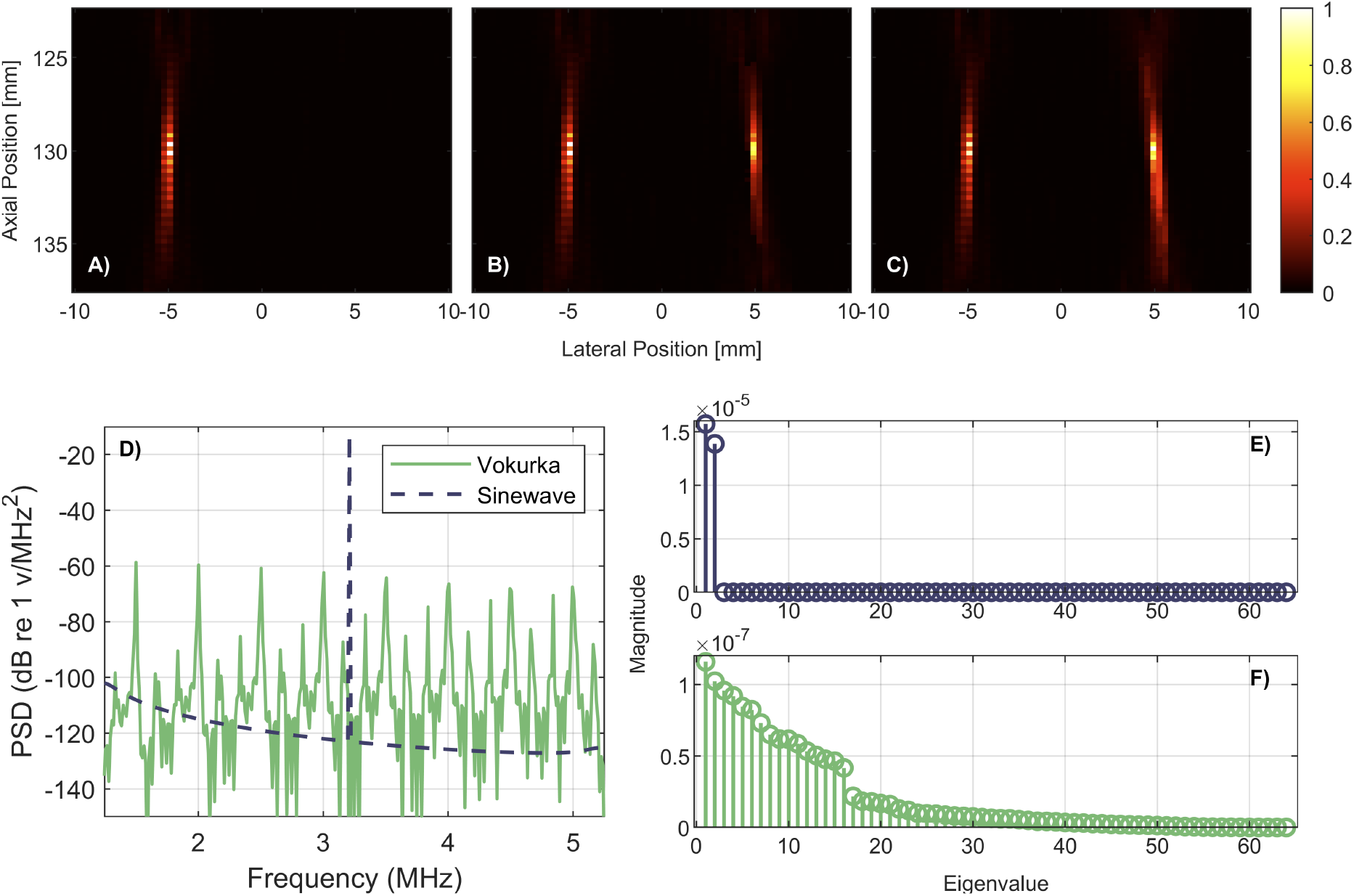
Panels A) - C) display PADAM images constructed with RF data where a sine-wave of amplitude 1 *MPa* and 3.21 *MHz* is positioned on the left, and a Vokurka source with mean peak amplitude 1 *MPa* is positioned on the right. Panel A) is beamformed with *m* = 2, panel B) shows *m* = 3, and C) shows *m* = 4 Panel D) shows the Eigenvalues of a sine-wave at the first pixel in the image, and panel E) shows the Eigenvalues of a Vokurka source at the same pixel. F) Shows the spectrum of the sine-wave and Vokurka signals in isolation.

Sinewave signals generate a spatial covariance matrix that has exactly two nonzero Eigenvalues, which are strong compared to the many nonzero Eigenvalues of the Vokurka source, seen in Figure 5E). At *m* = 2, only the sine-wave is visible in the entire image, as it is highly correlated and dominates the signal, even at the Vokurka source’s pixel location. However, at *m* = 3, more Eigenvalues are included than the sinewave’s alone, allowing one from the Vokurka source. The Vokurka source appears at approximately half the intensity of the sinewave source, since two Eigenvectors belong to the sinewave and only one from the Vokurka source. At *m* = 4, both regions appear similar in intensity. Beyond *m* = 4, more Eigenvectors correspond to the Vokurka source, shifting the visual weight primarily to Vokurka source. Notably, the Eigenvectors in PADAM are not scaled by their Eigenvalues, leading to a stacking effect. Admitting too many sources results in overlaying Eigenvectors from closely located sources, causing the 2 - 3 pixels to dominate the image. This can misidentify the strongest source’s location due to factors like the tail artifact, if *m* is set too high.

To further demonstrate this concept, four Vokurka sources were arranged in a rectangular configuration with one at the center. The time-condition of the Vokurka model was increased for the four corner sources, altering the pulse widths and increasing the ratio of harmonic to inharmonic energy. This serves as a pseudo-model for stable and inertial cavitation, where low-*θ* represents inertial cavitation and high-*θ* represents stable cavitation. Figure 6 presents three values of *m* from 1 to 3, showing that the central Vokurka-modeled inertial source has lower contrast at smaller *m*, which increases with *m*. Figure 6E illustrates the frequency characteristics of the two source models, confirming these observations: the largest two frequency domain peaks correspond to the stable sources, while the remaining peaks correspond to the inertial source.

**Figure 6:**
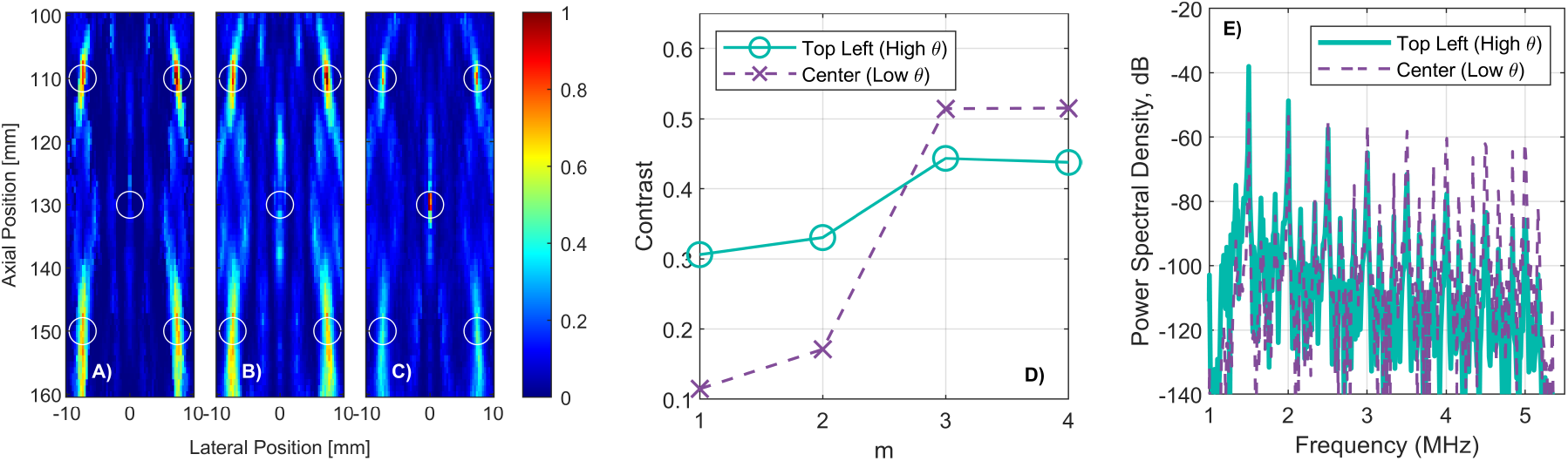
Panels A) Through C) show 3 PADAM images from *m*=1 to 3 for a model where 1 low-*θ* Vokurka model source (to mimic inertial cavitation) is placed in the center of the field of view, and 4 high-*θ* sources (to mimic stable cavitation) are placed at the four corners of the image. The high-*θ* source is suppressed by the larger frequency spectra peaks of the low-*θ* sources for *m*= 1 and 2 as seen in the contrast plot in panel D), because the two strongest peaks in the frequency-spectra come from the low-*θ* sources, as seen in panel E). For *m*=3 and 4, the center source has enough relatively strong frequency peaks to gain contrast in the image.

Consequently, the most significant Eigenvalues in the image come from the stable sources, and PADAM detects these until *m* = 3, at which point Eigenvalues from the inertial source begin to appear. The contrast of the sources as shown in Figure 6D highlights this effect. This finding suggests that PADAM may be used to distinguish between stable and inertial cavitation in an image, an aspect further explored in the in-vitro methodology of this work.

### In Vitro Models

To test if PADAM could provide superior artifact reduction in-vitro as seen in the computational models, we performed an in vitro experiment in a tube phantom. Figure 7 shows cavitation images beamformed with DSI and RCB (*ε* = 10) for comparison, and PADAM images with *m* = 1, 5, and 10. Among these, PADAM effectively suppresses reflections in the region, revealing the true cavitation source on the left side of the focal area.

**Figure 7:**
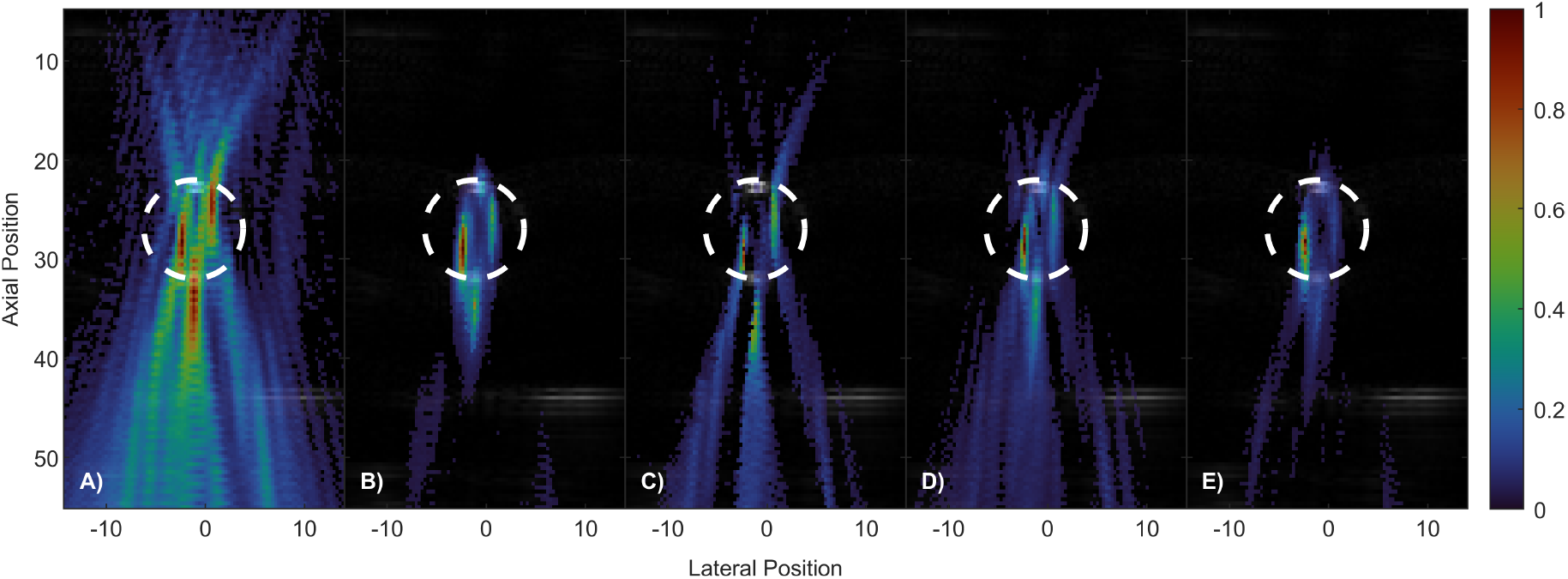
Panels A) and B) show passive cavitation images for the in-vitro model using DSI and RCB respectively, overlaid onto a B-Mode image of the same field-of view. A white dotted line is shown where the 2 mL tube’s boundary lies. The near and far edges of this tube may be seen faintly in the B-mode image. Panels C), D), and E) show the same images beamformed with PADAM, for *m*= 1, 5, and 10 respectively. PADAM demonstrates the superior artifact reduction.

PADAM cavitation images of the in-vitro skull model are shown in Figure 8. The images display significant improved localization capability of PADAM through a skull phantom in comparison with RCB and DSI. RCB displays a limitation where it was not able to remove most of the artifactual noise in the image, despite an extensive *ε* parameter search, although it did remove the tail artifact in comparison to DSI. DSI had an MSI of 0.015032 (up to 12.3% noise), in comparison to RCB with 0.022674 (up to 15.1% noise) and PADAM with 0.001486 (up to 3.8% noise). PADAM has the primary benefit of noise and artifact reduction in this case; the location of the cavitation center is similar in the three images, and matches the known location of the tube from the B-Mode image. PADAM localizes the source to slightly below the tubing, which is the expected effect of focal aberration. Interestingly, the aberration is easier to identify with PADAM because of its improved axial resolution. Axial and lateral beam profiles at the focal point of the cavitation image are included in the supplementary material.

**Figure 8:**
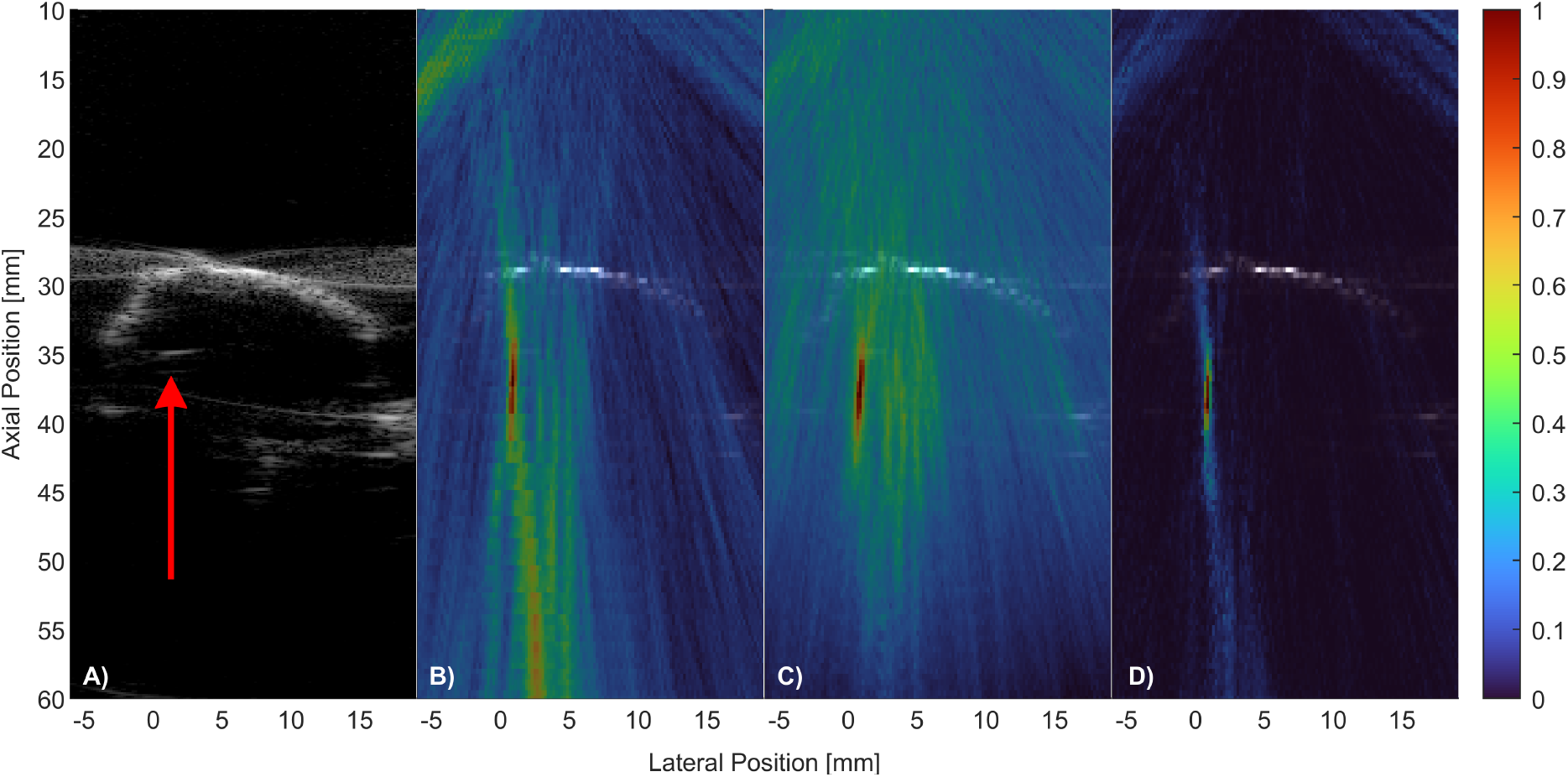
Passive Cavitation images from the rat skull phantom model. The rat skull is seen in white at a depth of 35 mm, indicated by the red arrow in the B-Mode image in panel A). Each panel shows the same frame of RF-Data beamformed with B) DSI, C) RCB, and D) PADAM, *m* = 4.

## 5 Discussion

Our results demonstrate that PADAM in the time domain is a highly effective approach for accurate passive cavitation imaging. PADAM consistently outperformed DSI and RCB in both lateral and axial resolution, as well as point-spread function size for the observed areas in the imaging plane. Furthermore, PADAM exhibited superior reduction in tail artifacts compared to RCB for both singular sources and clusters, both in-silico and in-vitro, with and without a skull.

A particularly notable advantage of PADAM is its parameter *m*, which quantifies the frequency richness of incoming signals. As previously discussed and shown in the supplementary material, sinewaves have exactly two nonzero Eigenvalues. It turns out that the exact number of nonzero Eigenvalues present in a signal’s spatial covariance matrix is 2 times the number of frequencies in that signal [33]. Figure 3E in the supplementary material also shows the Vokurka source’s Eigenvalue pattern, which has 16 Eigenvalues that represent the 8 in-band harmonics of the drive frequency, which was 500*KHz*, seen in panel D). In this way, PADAM, despite being a time-domain beamformer, can filter by frequency content. This provides a physically meaningful and intuitive measure that enables differentiation between frequency spectra and, consequently, between cavitation mechanisms.

As discussed in the Introduction, stable and inertial cavitation induce distinct bio-effects: stable cavitation facilitates targeted vascular permeation, whereas inertial cavitation enables tissue ablation. By adjusting *m*, users can localize sources or clusters (i.e., setting *m* to 1) or increase *m* beyond the number of sharp harmonics or ultraharmonics to visualize inertial cavitation regions. PADAM’s ability to dynamically distinguish cavitation mechanisms has significant implications in image-guided FUS therapy, particularly for applications requiring selective monitoring of stable versus inertial cavitation. This is crucial when avoiding unintended bio-effects from inertial cavitating bubbles in sensitive areas.

Compared to RCB, PADAM is easier to use and more computationally efficient. Unlike RCB, which requires parameter tuning (i.e., *ε*) to optimize performance, often resulting in inconsistent image quality, PADAM does not require exhaustive parameter searches, making it more userfriendly. Additionally, PADAM is computationally faster than RCB, as it relies on Eigenvalue decompositions rather than matrix inversions for each pixel. Furthermore, the algorithm’s flexible handling of signal and noise subspaces allows for rapid computation of multiple images with varying *m* values, rather than rerunning the algorithm separately for each case. PADAM is a candidate for parallelization and speedup by using algorithms that identify the largest several Eigenvalues (namely, the power method) instead of every Eigenvalue.

However, a key limitation of PADAM is that it is not a power-based beamformer, and should not be used to monitor cumulative energy absorption. As discussed in Section 3, PADAM assigns pixel values inversely proportional to noise correlation—essentially measuring what is not noise. Future work could explore using PADAM images as masks over a power-based beamformer to generate comprehensive dosage maps.

There were also several experimental limitations in both the in-silico and in-vitro methods. The Vokurka model, while useful, assumes a smooth ramp-down of the impulse pressure, which oversimplifies actual cavitation behavior. In addition, this study also varied the Vokurka *θ* time-condition parameter to mimic bubble dynamics. While altering *θ* modified the frequency spectrum and the ratio of harmonic to inharmonic energy, it did not generate the ultraharmonics commonly observed in cavitation behavior. Furthermore, the inharmonic content in the Vokurka model mainly arises from aliasing rather than a well-defined inertial cavitation mechanism, limiting its ability to fully replicate stable and inertial cavitation. However, the fundamental principle underlying PADAM, differentiating sources dynamically based on frequency content, remains effectively demonstrated and was further validated by the experimental results.

The experimental setup was designed to capture both inertial cavitation at the FUS focal center and stable cavitation in the surrounding periphery. Future studies could investigate bubbles in a thin-tube phantom at low concentrations to track individual bubble dynamics over time as they migrate into a FUS focal region. Another limitation was the skull model thickness. PADAM performed well in the presence of a thin rat skull at a low drive frequency; however, the wavelength of the FUS driving pulse is significantly longer than the thickness of the parietal skull fragment, at 0.75 mm. The pressure wave would have been minimally aberrated at this drive frequency, but any harmonics or ultraharmonics above 2 MHz will have a wavelength on the same order of magnitude as the skull thickness. Future studies could investigate the effect of primate skulls on PADAM performance. An in-vivo study could evaluate PADAM’s performance in a more realistic setting, and would provide insight into how PADAM differentiates between stable and inertial cavitation in a living organism. Lastly, extending PADAM to the frequency domain would enable direct frequency selection as it does with DSI.

## 6 Conclusion

This work introduces PADAM in the time domain as a passive cavitation imaging method and compares it to classic beamformers such as DSI and RCB. Our results demonstrate that PADAM offers significant advantages, including improved lateral and axial resolutions, artifact suppression, and a physically meaningful input parameter that is intuitive and requires minimal tuning. Additionally, PADAM is computationally more efficient than RCB, as it avoids matrix inversions for every pixel, offering a moderate speedup. Further work will focus on optimizing and parallelizing PADAM for clinical applications, as well as exploring its extension to the frequency domain to enhance its ability to differentiate between the stable and inertial cavitation modalities.

The capability of PADAM to distinguish cavitation mechanisms based on its input parameter represents a valuable advancement in passive cavitation imaging with broad implications. A potential clinical application could involve the use of PADAM as a mask on a DSI image to estimate the power of the signal while simultaneously identifying regions of stable and inertial cavitation. This dual capability can provide clinicians with critical insights for localizing bio-effects for targeted therapies, facilitating the advancement of ultrasound-guided focal therapies, and accelerating safe translation to patients.

